# Gut microbial disruption in critically ill patients with COVID-19 associated pulmonary aspergillosis

**DOI:** 10.1101/2022.10.08.511408

**Authors:** H. Carlo Maurer, David Schult, Plamena Koyumdzhieva, Sandra Reitmeier, Moritz Middelhoff, Sebastian Rasch, Markus List, Klaus-Peter Janssen, Katja Steiger, Ulrike Protzer, Roland M. Schmid, Klaus Neuhaus, Dirk Haller, Michael Quante, Tobias Lahmer

## Abstract

**Objectives:** COVID-19 disease can be exacerbated by *Aspergillus* superinfection (CAPA). The causes of CAPA are not yet fully understood. Recently, alterations in the gut microbiome have been associated with a complicating course and increasing severity of COVID-19 disease, most likely via immunological mechanisms. Aim of this study was to investigate a potential association between severe CAPA and alterations in the gut and bronchial microbiota.

**Methods:** We performed 16S rRNA gene amplicon sequencing of stool and bronchial samples from a total of 16 COVID-19 patients with CAPA and 26 patients without CAPA. All patients were admitted to the intensive care unit. Results were carefully tested for potential influences on the microbiome during hospitalization.

**Results:** We found that late in COVID-19 disease, CAPA patients exhibited a trend towards reduced gut microbial diversity. Furthermore, late stage CAPA disease showed an increased presence of *Staphylococcus epidermidis* in the gut. This is not found in late non-CAPA cases or early disease. The analysis of bronchial samples did not show significant results.

**Conclusions:** This is the first study showing alterations in the gut microbiome accompany severe CAPA and possibly influence the host’s immunological response. In particular, an increase of *Staphylococcus epidermidis* in the intestine could be of importance.

**Summary:** The composition of intestinal bacteria in severe CAPA disease is altered with an increase in Staphylococcus epidermidis in the gut. Alterations in the composition of intestinal bacteria in severe CAPA may indicate immunologic involvement of the gut in the disease.

## INTRODUCTION

Along with other uncertainties during intensive care unit (ICU) treatment of patients affected by coronavirus disease 2019 (COVID-19), super- and/or co-infections are well-known complications of severe acute respiratory syndrome coronavirus type 2 (SARS-CoV-2) pneumonia in critically ill patients (1–3).

Similar to severe influenza pneumonia, this new viral disease can also lead to the emergence of invasive pulmonary aspergillosis in patients admitted to the ICU (4, 5). The so called COVID-19-associated pulmonary aspergillosis (CAPA) is an opportunistic secondary infection, primarily affecting ICU patients with severe acute respiratory distress syndrome (ARDS) (5). Patients affected by COVID-19 associated ARDS and CAPA suffer from an increased morbidity and mortality compared to patients without CAPA and COVID-19 ARDS (6, 7). Besides immunologic mechanisms caused by SARS-CoV-2 infection, invasive mechanical ventilation, concomitant corticosteroid or antibody treatment, and older age, were identified as independent risk factors for an increased susceptibility to invasive fungal infections (6, 7).

Changes in the gut microbiome are known to have an impact on the immune response via various cross talks between intestinal bacteria and immune cells (8, 9). As presented, in recent studies not only an altered microbial composition of the gut but also certain individual bacteria may contribute to the inflammatory response and disease severity in COVID-19 patients (10–13). Recently, our group has shown that changes in intestinal bacterial composition are associated with an increased complication rate of COVID-19 (13). CAPA is a severe infectious complication of COVID-19, and building on our previous results, we addressed the importance of the gut on the development of CAPA in the same patient cohort.

## METHODS

### Study design and patient cohort

The aim of this single center observational study at the Klinikum rechts der Isar of the Technical University of Munich was to investigate possible associations between an altered intestinal and lung microbial composition and the occurrence of COVID-19-associated pulmonary aspergillosis (CAPA) in critically ill patients.

To this end, we examined a total of 42 patients treated in our ICU for COVID-19 induced respiratory or multi-organ failure, between April and July 2020. SARS-CoV-2 infection was confirmed by quantitative reverse transcription PCR (TaqMan™-PCR performed on Roche cobas® 6800, Basel, Switzerland), performed on nasopharyngeal swabs or tracheal fluid. During the ICU stay, patients were prospectively screened using respiratory specimens in defined time intervals (every three days during their ICU stay) for the development of CAPA. Beside galactomannan testing from serum and respiratory specimens, routine microbiological tests were performed at the same time intervals. All patients received a computed tomography (CT) scan of the chest before ICU admission. Sixteen patients (38%) were diagnosed with CAPA at the time of study enrollment, based on the ECMM/ISHAM consensus criteria (14). Every COVID-19 patient fulfilling the mentioned criteria was discussed between experts in microbiology, infectious disease and intensive care medicine to ascertain that the criteria for CAPA were met. The remaining 26 patients served as CAPA-negative COVID-19 controls. The severity of COVID-19 disease was defined using the WHO Ordinal Scale for Clinical Improvement in Hospitalized Patients with COVID-19 (15). All patients were seriously ill with an average WHO grade of 6.8 (standard deviation: 1.33), and some patients died in the course of the disease (N = 15). Patients were fed parenterally together with enteral nutrition via a gastric tube (artificial feeding). Antibiotic therapies were classified according to their effect on the gut microbiome as described previously (13). We evaluated the antibiotic intake up to two weeks before the stool or sputum sample was taken. No vancomycin or linezolid was administered.

### Microbial sampling and 16S rRNA gene sequencing

We collected a total of 14 tracheal and 40 stool samples for microbial profiling from the 42 COVID-19 patients treated in our ICU. Microbial samples had matching complete blood count (CBC) and serum samples available. For intubated patients, stool was collected either by a Flexi-Seal™ (Convatec, Munich, Germany) device (N = 13), a fecal collector (stool collection bag, N = 23) or regular means (N = 4). Tracheal secretions were obtained by endobronchial aspiration via respiratory tube or by bronchoalveolar lavage during intubation.

Preparation and sequencing of microbial samples was carried out as described previously (13). Samples were stored in a solution to stabilize DNA (MaGix PBI, Microbiomix GmbH, Regensburg, Germany). Sample preparation and paired-end sequencing was performed on an Illumina MiSeq targeting the V3V4 region of the 16S rRNA gene. Raw FASTQ files were processed using the NGSToolkit (https://github.com/TUM-Core-FacilityMicrobiome/ngstoolkit) based on USEARCH 1125 to generate denoised zero-radiation operational-taxonomic units (zOTUs). We excluded zOTUs with a relative abundance <0.1% and a prevalence <5%. Assessment of alpha diversity and taxonomic binning were computed using the Rhea software pipeline (16).

### Statistical analysis

Statistical analysis of 16S rRNA profiles was conducted as described previously (13). Briefly, read counts were normalized and differences in relative abundance of taxa and/or zOTUs were determined by a Kruskal-Wallis Rank Sum test for multiple groups and Mann-Whitney-U tests for pairwise comparisons, respectively. Associations between categorical variables were tested using Fisher’s Exact test. Similarity between samples was estimated using on a distance matrix based on generalized UniFrac distances. Confounders and possible effect modifiers of microbial ecosystems were determined via a permutational multivariate analysis of variance using the aforementioned distance matrix.

### Ethical approval

Patients signed the informed consent either before intubation or after extubation. For patients who have died, the ethics committee permits the use of the anonymized data. The institutional review board for human studies approved the protocols and written consent was obtained from the subjects or their surrogates if required by the institutional review board. The study was conducted in accordance with the declaration of Helsinki and approved by the ethics committee of the Technical University Hospital of Munich (221/20 S-SR).

## RESULTS

### CAPA occurs more frequently among elderly COVID-19 patients and indicates an unfavorable clinical course

Among our cohort of 42 COVID-19 patients selected for microbial sampling, 16 (38%) received a diagnosis of CAPA (**Figure 1A**). A basic summary of clinical characteristics and patient-level information are provided in **Supplementary Tables 1 and 2**. We first examined clinical covariates for their association with CAPA status and only found age to differ significantly between groups. Specifically, the median age among COVID-19 patients with CAPA was 76.5 years as opposed to 61.5 years in non-CAPA patients (**Figure 1B**, p = 0.008, Mann-Whitney-U test). Further trends of a higher occurrence in CAPA patients could be appreciated for age-related pre-existing conditions such as coronary artery disease and atrial fibrillation (**Suppl. Table 1 and Suppl. Figure 1A**). Consistent with previous reports (17) COVID-19 patients with pulmonary *Aspergillus* superinfection exhibited a worse course of disease, prolonged mechanical ventilation (**Suppl. Table 3 and Suppl. Figure 1B**) and higher fatality rates (**Figure 1B**). Accordingly, vasopressor therapy was needed significantly more often in CAPA patients (**Figure 1C**). Importantly, the application of dexamethasone was balanced between CAPA and non-CAPA cases (**Suppl. Table 3**). Taken together, we confirmed that CAPA is associated with a significantly increased COVID-19 disease severity, and we found older age to be the main risk factor for contracting CAPA in our cohort.

**Figure 1.**
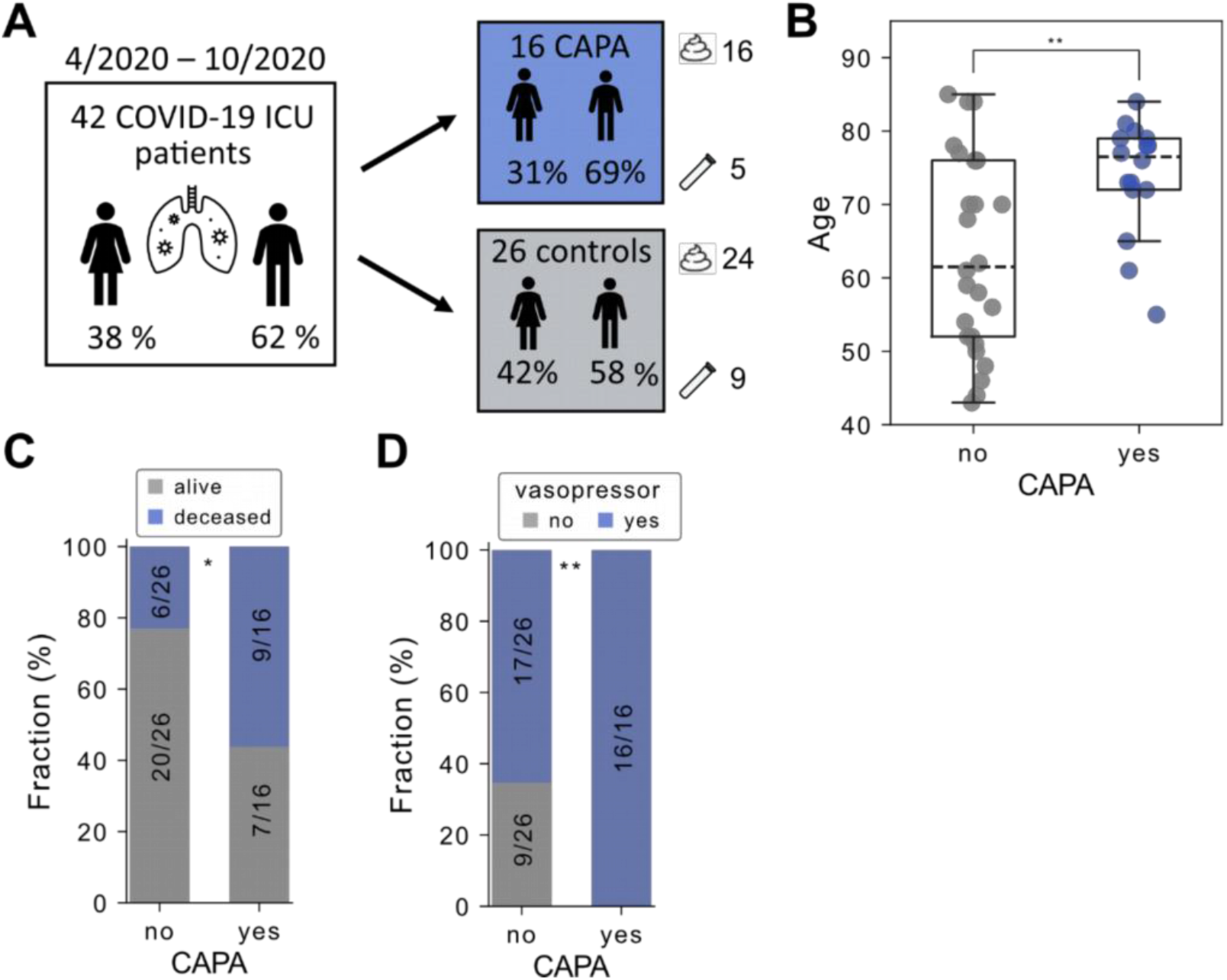
(**A**) Overview of the study design. Stool and tracheal secretions are indicated. (**B**) Distribution of age according to COVID-19-associated pulmonary aspergillosis (CAPA). P-value derived from a twotailed Mann-WhitneyU test. (**C**) Fraction of deceased patients according to CAPA status. (**D**) Fraction of patients requiring vasopressor therapy according to CAPA status. In boxplots, the box ranges from Q1 (the first quartile) to Q3 (the third quartile) of the distribution and the range represents the IQR (interquartile range). The median is indicated by a dashed line across the box. The “whiskers” on box plots extend from Q1 and Q3 to 1.5 the IQR. Unless otherwise specified, p-values are derived from two-tailed Fisher’s exact test. *** p <= 0.001; ** p <= 0.01; * p <= 0.05; ns, not significant.

### Consideration of factors influencing the microbiome and comparability between study groups

16S rRNA gene amplicon sequencing of 40 fecal samples available from the 42 COVID-19 patients treated in our ICU revealed diverse ecosystems dominated by the two major phyla *Firmicutes* and *Bacteroidota* (cumulative mean relative abundance, 85%). Importantly, we did not detect significant differences in gut microbial composition between CAPA and non-CAPA patients (**Figure 2A and Suppl. Figure 2B**). Consequently, we evaluated further sample and patient metadata more carefully in an unbiased manner using a multivariate analysis of inter-individual variabilities in microbiota structure (**Suppl. Table 4**). Here, we found ‘days since hospital admission’ as the most prominent factor (p = 0.001) (**Figure 2B**). Both, species richness and the abundance of a total of 47 genera correlated significantly (false discovery rate < 0.1) with days passed since hospital admission (**Suppl. Figure 2A**). Most genera showed a negative relationship, including *Actinomyces, Ruminococcus* and *Bifidobacterium*. On the other hand, *Enteroccocus* and *Staphylococcus* genera represented exceptions with increasing relative abundance during hospital stay.

**Figure 2.**
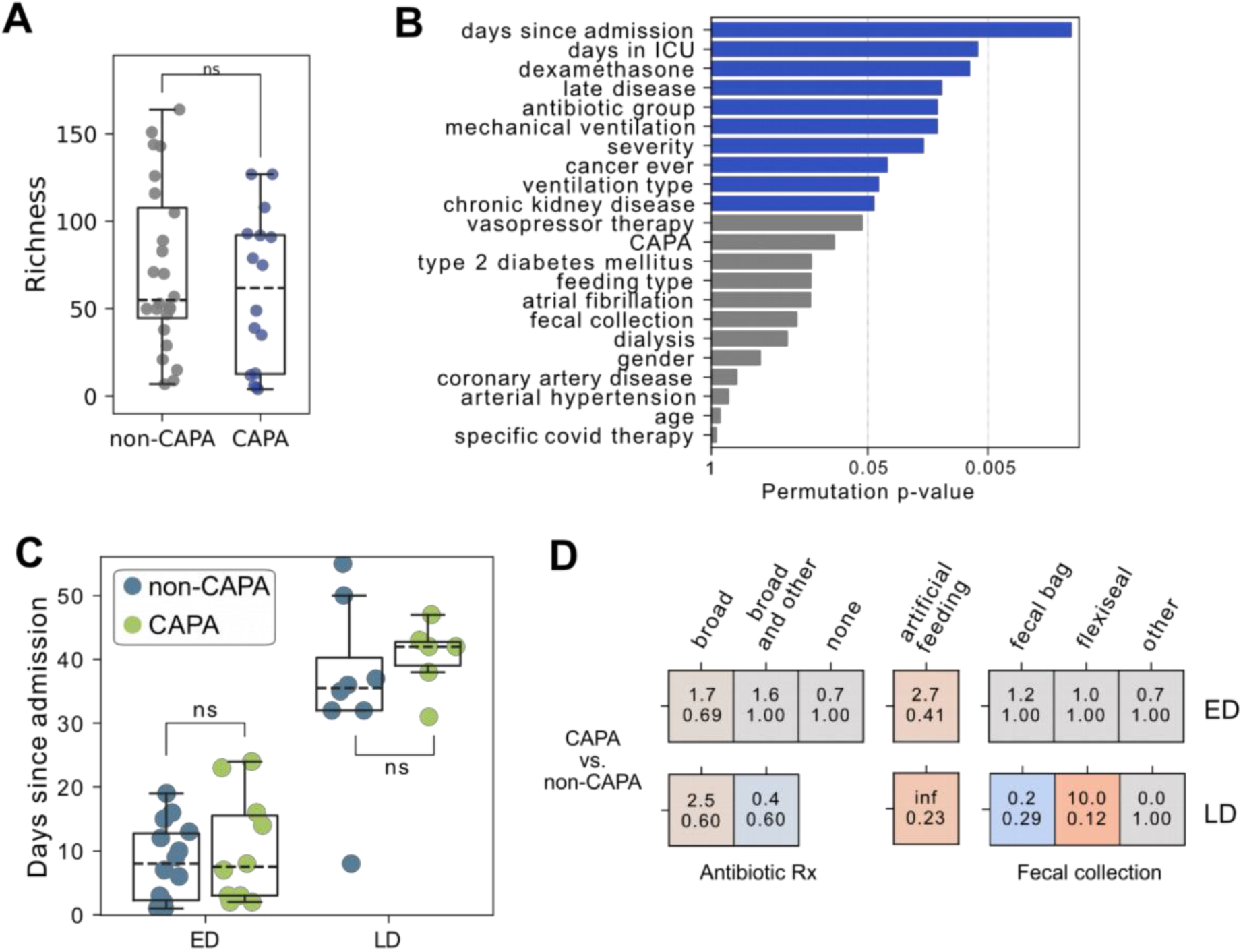
(**A**) Richness (=number of detected zOTU) on y-axis as a measure of sample diversity across stool samples from COVID-19 with and without CAPA. (**B**) P-values derived from a permutational multivariate analysis of variances (x-axis) for the indicated clinical covariates and the stool microbial ecosystem diversity as represented by generalized UniFrac distance. Blue bars mark significant covariates. (**C**) Days since admission to the hospital (y-axis) across patients with early vs. late COVID-19 disease and with or without Aspergillus superinfection (**D**) Association of antibiotic treatments, feeding type and stool collection method (x-axis) and early and late COVID-19 disease with Aspergillus infection (y-axis). For each combination and cell, Odds ratio (OR, upper number) and p-value (from a two-sided Fisher’s exact test) are depicted. Cells are colored by p-value and the sign of log(OR). In boxplots, the box ranges from Q1 (the first quartile) to Q3 (the third quartile) of the distribution and the range represents the IQR (interquartile range). The median is indicated by a dashed line across the box. The “whiskers” on box plots extend from Q1 and Q3 to 1.5 the IQR. Unless otherwise specified, p-values are derived from two-tailed Mann-Whitney-U tests. *** p <= 0.001; ** p <= 0.01; * p <= 0.05; ns, not significant.

Further influential covariates included ‘total length of ICU stay’, dexamethasone and antibiotic treatment, and whether patients were considered to be late in their COVID-19 disease course, i.e. if they had already cleared the virus from their upper airway as verified by PCR testing (**Figure 2B**). In contrast to the above covariates, we did not find feeding, stool collection type or age to exhibit significant influence on the global microbiota structure in these 40 stool samples.

Patients with late COVID-19 were both hospitalized and treated in the ICU for a significantly longer time than their early disease counterparts (**Suppl. Figure 2C, D**). Furthermore, dexamethasone treatment was absent in late disease patients and fecal management more often involved a Flexi-Seal™ device (**Suppl. Figure 2E**) and a lack of antibiotic therapy tended to occur more in patients with early COVID-19 disease. Lastly, stool microbial alpha diversity was significantly lower in patients with late COVID-19 disease (**Suppl. Figure 2F**).

For further investigation into microbial changes accompanying CAPA, we accounted for the aforementioned factors by stratifying patients both by early (ED) versus late disease (LD) and non-CAPA versus CAPA status yielding four groups: ED non-CAPA (N = 15), ED CAPA (N = 10), LD non-CAPA (N = 9) and LD CAPA (N = 6). With this approach, no significant difference in hospitalization length (**Figure 2C**), antibiotic therapy, feeding or stool collection method could be observed between the respective CAPA and non-CAPA groups (**Figure 2D**), allowing us to draw firm conclusions on microbial composition.

We further examined microbial samples from tracheal secretions, which were available from 9 non-CAPA and 5 CAPA patients (data not shown). Permutation analysis of inter-individual variabilities in microbiota structure found no significant role for CAPA status and again identified ‘days since admission’ as the most influential covariate. The latter was highest in four late disease COVID-19 patients. However, the limited patient number in this subset prohibited further statistically sound comparisons between non-CAPA and CAPA patients.

In summary, stool and bronchial microbial communities showed increased damage with hospital stay. Adjustment for SARS-CoV-2 disease course status yielded groups of CAPA and non-CAPA patients that were balanced concerning major influences on stool inter-individual variabilities.

### Impaired fecal microbiota and staphylococcal outgrowth occurs in late but not early CAPA

In the following, we analyzed the association CAPA and gut microbial signatures in more detail. In the ED group, no difference in alpha diversity could be demonstrated for CAPA or non-CAPA patients (p = 0.89, Mann-Whitney-U test). On the other hand, in the group of LD, patients with CAPA tended to show a decrease in microbial richness compared to non-CAPA cases, although this was not significant according to the p-value (p = 0.11, **Figure 3A**). Similarly, LD CAPA samples were significantly overrepresented (Odds ratio = 15.0, p = 0.01 Fisher’s Exact test) among the least diverse 20 percent of samples as determined by sample richness (**Figure 3B**). No linear correlation could be observed for changes in gut microbial composition between CAPA and non-CAPA patients in ED versus LD respectively (**Figure 3C**), suggesting that pulmonary aspergillosis associates with distinct metagenomic shifts in each condition. Interestingly, significantly more *Staphylococcus epidermidis* was found in the stool of LD CAPA patients compared with LD non-CAPA patients. (p: 0.008, **Figure 3D**). Thus, we observed both a trend towards a reduced gut microbial within-sample diversity and a specific expansion in *S. epidermidis* relative abundance separating CAPA from non-CAPA cases late in COVID-19 disease.

**Figure 3.**
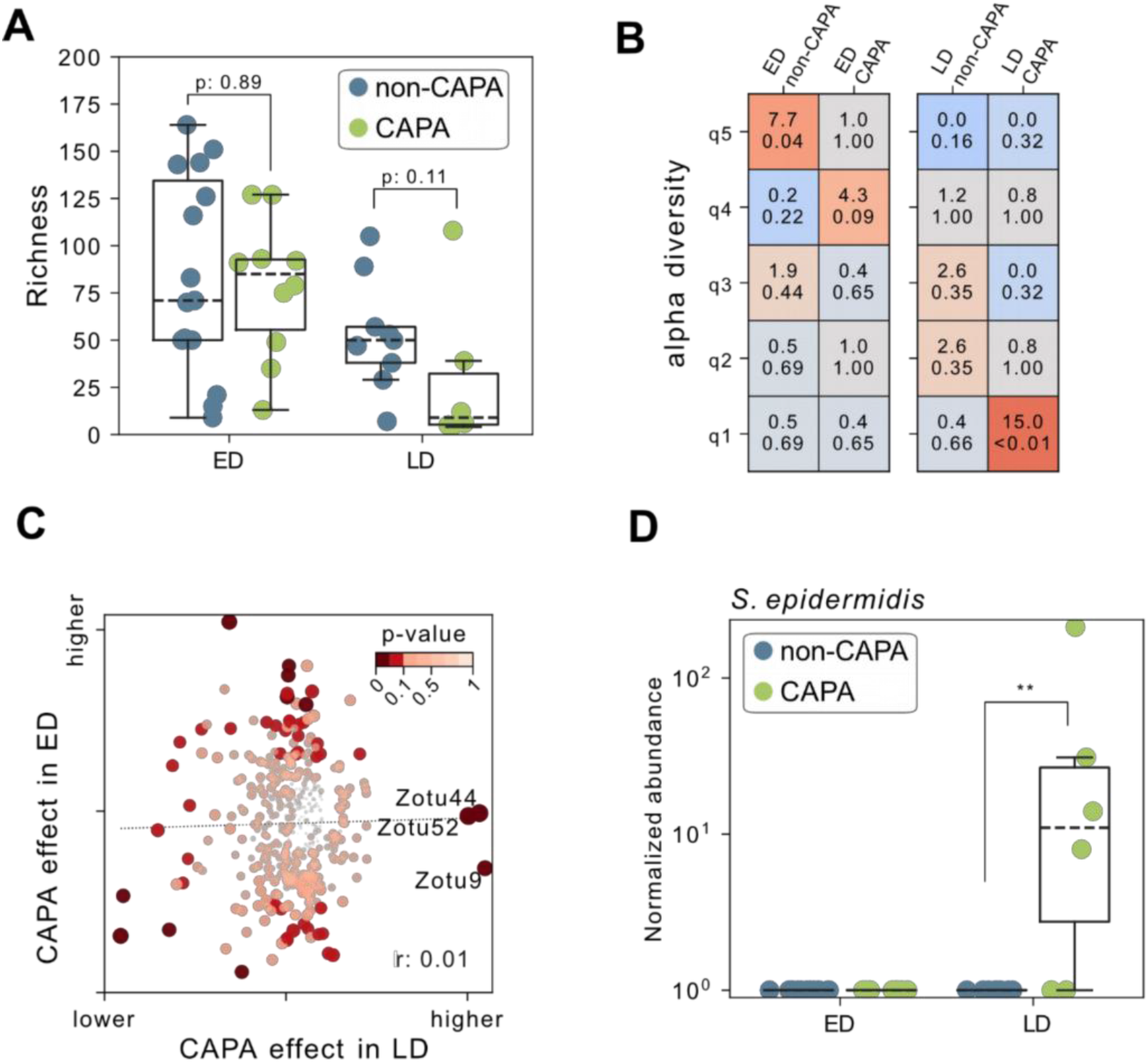
(**A**) Richness (= number of detected zOTU) on y-axis across patients grouped according to COVD-19 disease stage (ED, early disease vs. LD, late disease) and Aspergillus superinfection. (**B**) Association of microbial diversity quintile (q1 = lowest to q5 = highest) (y-axis) and patient groups according to early (ED) vs. late (LD) COVID-19 infection and presence of CAPA. For each combination and cell, Odds ratio (OR, upper number) and p-value (from a two-sided Fisher’s exact test) are depicted. Cells are colored by p-value and sign of log(OR). (**C**) Scatter plot illustrating the effect of Aspergillus superinfection on microbial signatures (= difference between U2 and U1 from a Mann-Whitney-U test) for stool specimen from late disease (x-axis) and early disease (y-axis) patients, respectively. Dots (= individual zOTU) are colored by the minimum p-value from both comparisons as calculated from a twosided Mann-Whitney-U test. (**D**) Median normalized abundance on y-axis from zOTUs belonging to S. epidermidis across patients grouped according to COVID-19 disease stage and Aspergillus superinfection. In boxplots, the box ranges from Q1 (the first quartile) to Q3 (the third quartile) of the distribution and the range represents the IQR (interquartile range). The median is indicated by a dashed line across the box. The “whiskers” on box plots extend from Q1 and Q3 to 1.5 the IQR. Unless otherwise specified, p-values are derived from two-tailed Mann-Whitney-U tests. *** p <= 0.001; ** p <= 0.01; * p <= 0.05; ns, not significant.

## DISCUSSION

COVID-19 has now been a global pandemic since late 2019, impacting many aspects of life. Although vaccines have reduced morbidity, COVID-19 remains a threatening and serious disease, especially in predisposed and immunocompromised patients. Viral infections of the lung are prone to fungal superinfection (18) likely facilitated by inflammatory damage to the lung epithelium and an impaired immune response associated with the disease (18, 19). Coinfection, on the other hand, leads to a significant worsening of respiratory symptoms with increased mortality in patients with viral pulmonary infections (20). This also applies to SARS-CoV-2 infection, in which coinfection with *Aspergillus fumigatus*, can lead to COVID-19-associated pulmonary aspergillosis with poor outcome and the need for intensive care (5, 7). Risk factors of CAPA disease include severe lung injury in the setting of COVID-19, older age, and preexisting pulmonary disease (21). In addition, the use of broad-spectrum antibiotics has been associated with the development of invasive pulmonary aspergillosis (IPA) (22), and the use of azithromycin also appears to predispose to CAPA (23). While systemic corticosteroids are known risk factors for IPA (24), data on their role in increasing the risk for CAPA remain controversial (7, 21)

The gut microbiome has critical influence on the maturation of intestinal lymphoid immune defenses (25) and, moreover, on the immune response in peripheral tissues via multiple cross talks with immune cells (9, 26, 27). There is evidence of interactions between intestinal bacteria and mycobiota (28) and bacterial metabolites such as short chain fatty acids seem to play a role in this inter-kingdom communication (29). Studies in mice have shown that intestinal microbiota influence the adaptive immune response to *Aspergillus fumigatus* in the lung via regulation of CD4 cells (30). A disturbed gut microbial balance between bacteria and fungi, in combination with an impaired immune response, can potentially contribute to invasive aspergillosis as described in the context of IPA and CAPA (31), but studies on the relationship between CAPA and the gut microbiome are lacking.

Here, we were able to show for the first time that CAPA patients exhibit impaired stool and bronchial microbial communities late in COVID-19 disease. Specifically, this includes a trend towards reduced species richness and a significant relative increase of *S. epidermidis* in fecal samples. We note that the limited number of cases for now preclude firm conclusions and further studies with higher numbers of cases are needed.

*S. epidermidis* belongs to the coagulase-negative staphylococci and is a facultative pathogen, managed for its high rate of nosocomial catheter and bloodstream infections. The bacterium furthermore serves as a genetic reservoir for *S. aureus*, increasing the pathogenicity and antibiotic resistance of *S. aureus* via gene transfer (32). While *S. epidermidis* occurs mainly on the skin of adults, the bacterium is increasingly found in the stool of preterm infants (33). Studies on the prevalence and function of *S. epidermidis* in the adult gut are scarce. It has been shown that strains of *S. epidermidis* from the gut or skin do not differ essentially in their ability to form biofilms and pathogenicity (34). A recent study has identified the gut as a reservoir and origin of bloodstream infection with *S. epidermidis*, thus challenging the informal dogma that this infection originates exclusively from the environment or the skin (35). In addition, colonization of the intestine with *S. epidermidis* in mice leads to serious damage of various extraintestinal organs, such as the kidneys and the liver (36).

To consider potential confounders on the microbiome in the context of hospitalization, we addressed antibiotic therapy, diet, length of hospital stay, and method of stool collection. These factors were balanced in the late COVID-19 disease, which allowed for reasonable comparability between non-CAPA and CAPA cases. Nevertheless, we cannot rule out conclusively, that the microbial results were influenced by the circumstances of hospitalization and further functional studies are needed. Moreover, all samples were collected during the early phase of the pandemic when the SARS-CoV-2 wild type strain dominated. The influence of different virus strains on the composition of gut microbiota and development of CAPA need further investigation.

Nevertheless, our findings of an increase in fecal *S. epidermidis* in combination with the tendency of decreased microbial within-sample diversity in CAPA versus non-CAPA patients late in their COVID-19 disease may be indicative of an impaired gut microbiome in severe CAPA. The immunological consequences of the high percentage of coagulase-negative staphylococci the stool of patients with late CAPA disease are unclear. However, due to the option of addressing these changes by application of probiotics or adaption of stool management during hospitalization, the finding may be of therapeutic relevance in this severe disease. Further studies including these therapeutic options and larger case numbers are needed for validation.

## Acknowledgments

We thank all healthcare workers who made this study possible through their daily efforts in the ICU within the pandemic. We are grateful to the team around Dr. Christoph Spinner and Dr. Johanna Erber. We would also like to thank Julia Horstmann and Lisa Fricke for their help in sample recruitment and we are grateful to Angela Sachsenhauser, Caroline Ziegler and Lukas Mix from the Core Facility Microbiome of the ZIEL Institute for Food & Health for excellent technical support in 16S rRNA gene sequencing.

## Notes

**Disclosure statement:** No potential conflict of interest was reported by the author(s).

### Competing Interest Statement

The authors have declared no competing interest.

