## Supplemental Figure 1 for "Gut microbial disruption in critically ill patients with COVID-19 associated pulmonary aspergillosis"

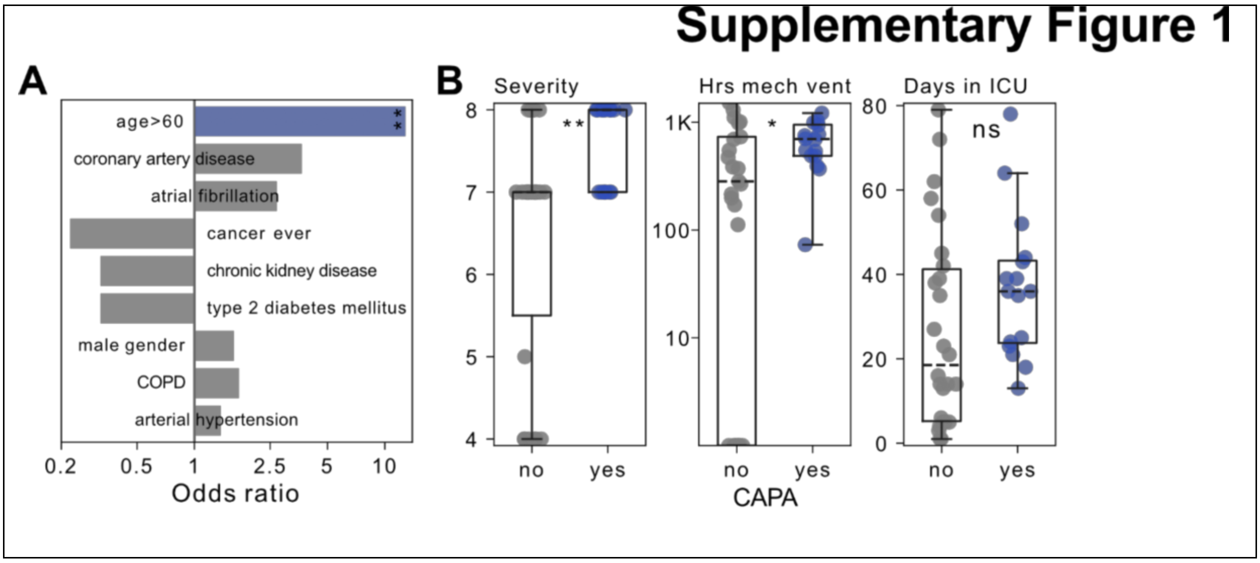


**Supplementary Figure 1.** (**A**) Odds ratios (x-axis) for the indicated clinical covariates reflecting their cooccurence with COVID-associated pulmonary aspergillosis (CAPA). Blue bars indicate covariates with significant results as determined from a two-sided Fisher's exact test. (**B**) WHO disease severity, hours of mechanical ventilation and days treated in an ICU according to CAPA status.

In boxplots, the box ranges from Q1 (the first quartile) to Q3 (the third quartile) of the distribution and the range represents the IQR (interquartile range). The median is indicated by a dashed line across the box. The “whiskers” on box plots extend from Q1 and Q3 to 1.5 the IQR. Unless otherwise specified, p-values are derived from two-tailed Mann-Whitney-U tests. *** p <= 0.001; ** p <= 0.01; * p <= 0.05; ns, not significant.
