## Supplemental Figure 2 for "Gut microbial disruption in critically ill patients with COVID-19 associated pulmonary aspergillosis"

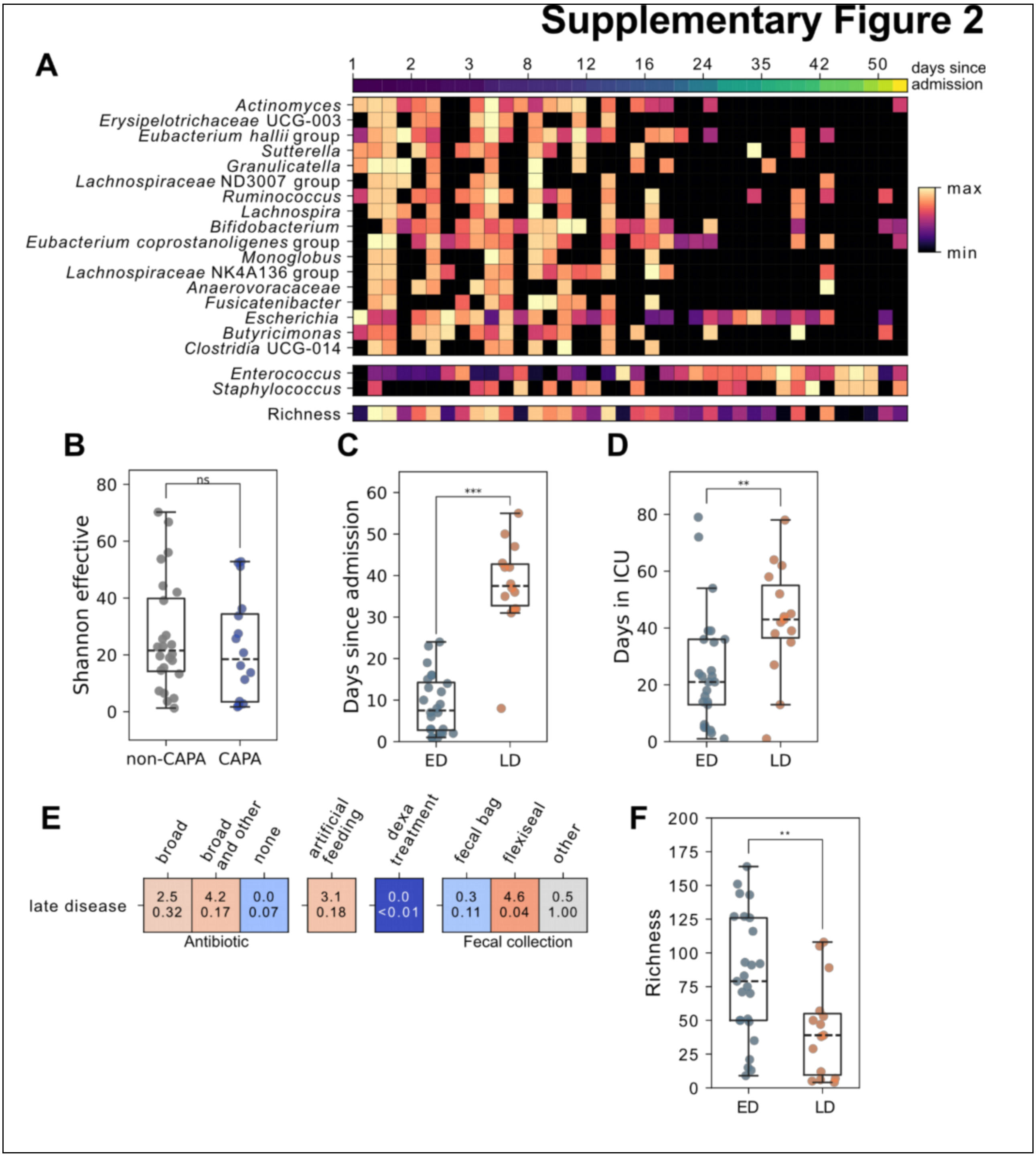


**Supplementary Figure 2.** (**A**) Heatmap representing quantile normalized abundance measures for the indicated genera (y-axis) and species richness ordered according to the time passed since hospital admission (xaxis). (**B**) Shannon effective number as a measure of alpha diversity among stool samples from COVID-19 patients with or without CAPA (**C**) Days since admission to the hospital according to COVID-19 disease stage (ED, early disease vs. LD, late disease). (**D**) Days spent in the ICU for patients with early and late COVID-19 disease (ED, early disease vs. LD, late disease) (**E**) Association of antibiotic treatments, feeding type and stool collection method (x-axis) and late COVID19 disease. For each combination and cell, Odds ratio (OR, upper number) and p-value (from a two-sided Fisher's exact test) are depicted. Cells are colored by p-value and the sign of log(OR). (**F**) Richness among microbial stool samples in early vs. late COVID-19 disease.

In boxplots, the box ranges from Q1 (the first quartile) to Q3 (the third quartile) of the distribution and the range represents the IQR (interquartile range). The median is indicated by a dashed line across the box. The “whiskers” on box plots extend from Q1 and Q3 to 1.5 the IQR. Unless otherwise specified, p-values are derived from two-tailed Mann-Whitney-U tests. *** p <= 0.001; ** p <= 0.01; * p <= 0.05; ns, not significant.
